## Supplementary material for "Extended-spectrum beta-lactamase (ESBL)-producing and non-ESBL-producing *Escherichia coli* isolates causing bacteremia in the Netherlands (2014 – 2016) differ in clonal distribution, antimicrobial resistance gene and virulence gene content": S1 Appendix

**EPIGENEC STUDY - SUPPORTING INFORMATION**

### S1 Appendix - content

**S1A Figure.** ST distribution among different onset of ECB

**S1B Table.** ST distribution among different onset of ECB

**S1C Figure.** ST distribution among different primary foci of ECB

**S1D Table.** ST distribution among different primary foci of ECB

**S1A Figure.** ST distribution among different onset of ECB[a]

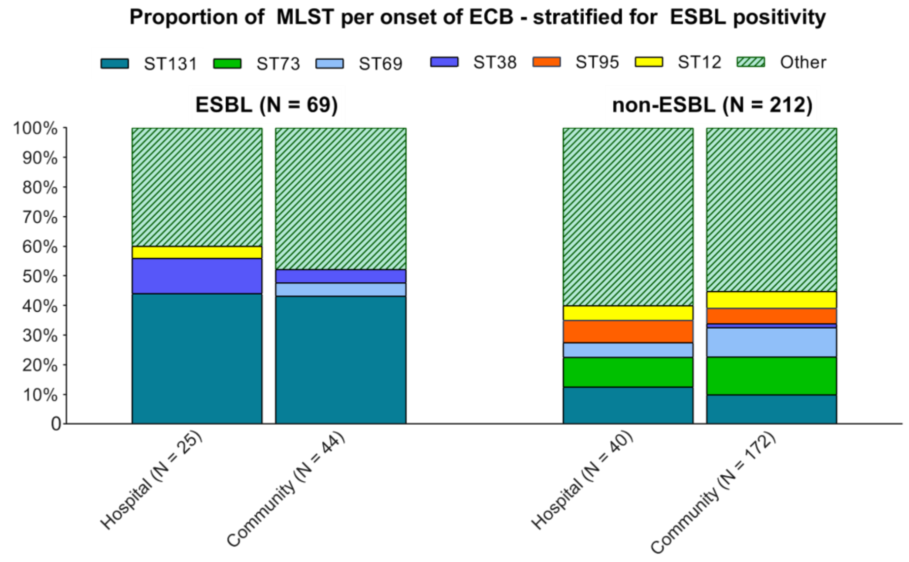


[a]ESBL-positivity based on phenotypic ESBL production.
ECB, *E. coli* bacteremia; ESBL, extended spectrum beta-lactamase; ST, sequence type.
Only STs that occurred >5% within non-ESBL-Ec or ESBL-Ec were grouped into main ST groups, the rest is categorized as "Other".

**S2 Table.** ST distribution among different onset of ECB

| | **ESBL *E. coli* [a]** | | **Non-ESBL *E. coli* [a]** | |
|---|---|---|---|---|
| | Hospital (N=25) | Community (N=44) | Hospital (N = 40) | Community (N = 172) |
| **ST131,** N (%) | 11 (44) | 19 (43) | 5 (13) | 17 (10) |
| **ST73,** N (%) | - | - | 4 (10) | 22 (13) |
| **ST69,** N (%) | - | 2 (5) | 2 (5) | 17 (10) |
| **ST38,** N (%) | 3 (12) | 2 (5) | - | 2 (1) |
| **ST95,** N (%) | - | - | 3 (8) | 9 (5) |
| **ST12,** N (%) | 1 (4) | - | 2 (5) | 10 (6) |
| **Other ST,** N (%) | 10 (40) | 21 (48) | 24 (60) | 95 (55) |

[a]ESBL-positivity based on phenotypic ESBL production.
ECB, *E. coli* bacteremia; ESBL, extended spectrum beta-lactamase; HB, hepatic-biliary; GI, gastro-intestinal; ST, sequence type

**S3 Figure.** ST distribution among different primary foci of ECB[a]

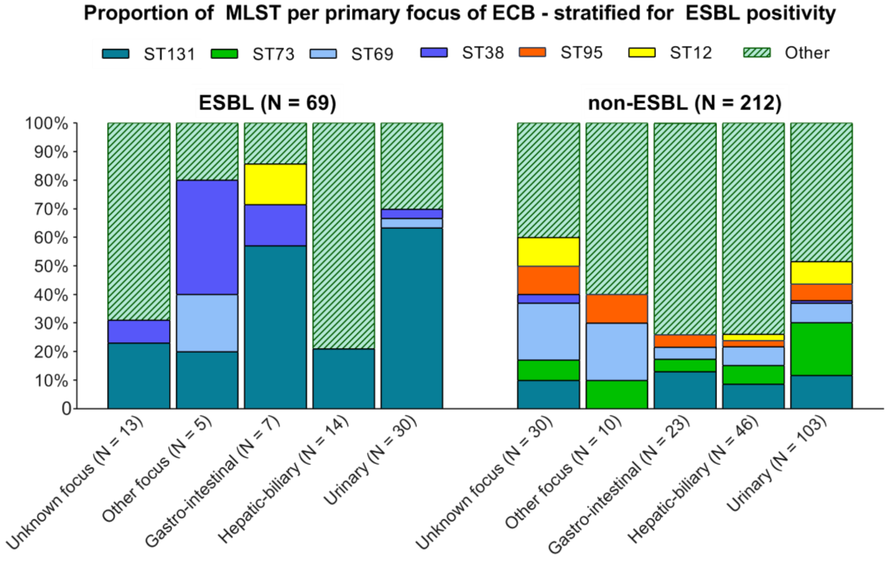


[a]ESBL-positivity based on phenotypic ESBL production.
ECB, *E. coli* bacteremia; ESBL, extended spectrum beta-lactamase; ST, sequence type
Only STs that occurred >5% within non-ESBL-Ec or ESBL-Ec were grouped into main ST groups, the rest is categorized as “Other”.

**S4 Table.** ST distribution among different primary foci of ECB[a]

| | **ESBL *E. coli*** | | | | | **Non-ESBL *E. coli*** | | | | |
|---|---|---|---|---|---|---|---|---|---|---|
| | Urinary (N=30) | HB (N=14) | GI (N=7) | Other (N=5) | Unknown (N=13) | Urinary (N=103) | HB (N=46) | GI (N=23) | Other (N=10) | Unknown (N=30) |
| **ST131,** N (%) | 19 (63) | 3 (21) | 4 (57) | 1 (20) | 3 (23) | 12 (12) | 4 (9) | 3 (13) | - | 3 (10) |
| **ST73,** N (%) | - | - | - | - | - | 19 (18) | 3 (7) | 1 (4) | 1 (10) | 2 (7) |
| **ST69,** N (%) | 1 (3) | - | - | 1 (20) | - | 7 (7) | 3 (7) | 1 (4) | 2 (20) | 6 (20) |
| **ST38,** N (%) | 1 (3) | - | 1 (14) | 2 (40) | 1 (8) | 1 (1) | - | - | - | 1 (3) |
| **ST95,** N (%) | - | - | - | - | - | 6 (6) | 1 (2) | 1 (4) | 1 (10) | 3 (10) |
| **ST12,** N (%) | - | - | 1 (14) | - | - | 8 (8) | 1 (2) | - | - | 3 (10) |
| **Other ST,** N (%) | 9 (30) | 11 (79) | 1 (14) | 1 (20) | 9 (69) | 50 (49) | 34 (74) | 17 (74) | 6 (60) | 12 (40) |

[a]ESBL-positivity based on phenotypic ESBL production.
ECB, *E. coli* bacteremia; ESBL, extended spectrum beta-lactamase; HB, hepatic-biliary; GI, gastro-intestinal; ST, sequence type
