## Supplementary material for "Extended-spectrum beta-lactamase (ESBL)-producing and non-ESBL-producing *Escherichia coli* isolates causing bacteremia in the Netherlands (2014 – 2016) differ in clonal distribution, antimicrobial resistance gene and virulence gene content": S2 Appendix

**EPIGENEC STUDY - SUPPORTING INFORMATION**

### S2 Appendix

**S2 Table.** Frequencies of O:serotypes per primary focus of ECB

| | **ESBL *E. coli* (N = 69)[a]** | | | | | **Non-ESBL *E. coli* (N = 212)[a]** | | | | |
|---|---|---|---|---|---|---|---|---|---|---|
| | Urinary (N=30) | HB (N=14) | GI (N=7) | Other (N=5) | Unknown (N=13) | Urinary (N=103) | HB (N=46) | GI (N=23) | Other (N=10) | Unknown (N=30) |
| **O25,** N (%) | 17 (57) | 2 (14) | 3 (43) | 1 (20) | 1 (8) | 12 (12) | 7 (15) | 2 (9) | - | 3 (10) |
| **O6,** N (%) | - | - | - | - | - | 15 (15) | 6 (13) | - | 2 (20) | 2 (7) |
| **O4,** N (%) | - | - | 1 (14) | - | - | 8 (8) | 1 (2) | - | - | 3 (10) |
| **O2/O50,** N (%) | - | - | - | - | - | 10 (10) | 6 (13) | - | 1 (10) | 2 (7) |
| **O75,** N (%) | - | - | - | - | - | 4 (4) | 1 (2) | 2 (9) | 1 (10) | 3 (10) |
| **O1,** N (%) | 1 (3) | - | - | - | - | 2 (2) | 1 (2) | - | - | 1 (3) |
| **O8,** N (%) | - | 4 (29) | - | - | - | 5 (5) | 3 (7) | 6 (26) | 2 (20) | 1 (3) |
| **O15,** N (%) | 1 (3) | - | - | 1 (20) | - | 5 (5) | - | - | - | 4 (13) |
| **O18,** N (%) | - | - | - | - | - | 5 (5) | - | - | 1 (10) | 1 (3) |
| **O16,** N (%) | 2 (6) | 1 (7) | 1 (14) | - | - | 2 (2) | - | 2 (9) | - | - |
| **Other serotype** | 9 (30) | 7 (50) | 2 (29) | 3 (60) | 12 (92) | 35 (34) | 21 (46) | 11 (48) | 3 (30) | 10 (33) |

[a]ESBL-positivity based on phenotypic ESBL production.

ESBL, extended spectrum beta-lactamase; HB, hepatic-biliary; GI, gastro-intestinal

Frequencies of all serotypes of the 4-valent and new potential 10-valent ExPEC vaccine are reported, the rest (including missing / unknown serotypes) is grouped as “Other serotype”. Percentages are column percentages.
