## Supplementary material for "Extended-spectrum beta-lactamase (ESBL)-producing and non-ESBL-producing *Escherichia coli* isolates causing bacteremia in the Netherlands (2014 – 2016) differ in clonal distribution, antimicrobial resistance gene and virulence gene content": S3 Appendix

### EPIGENEC STUDY - SUPPORTING INFORMATION

##### S3 Appendix - content

**S3A Table**. Detected acquired resistance genes with ResFinder 3.1.0 per antibiotic group

**S3B Figure.** Resistance gene count among epidemiological subgroups

**S3C Table.** Pairwise comparisons resistance gene score between epidemiological subgroups

**S3D Table.** Pairwise comparisons resistance gene score between STs

**S3A Table.** Detected resistance genes with ResFinder 3.1.0 per antibiotic group

| (Broad-spectrum) Beta-lactamases | | | ESBL and ampC | | | Macrolides | | | Fluoroquinolones | | |
|---|---|---|---|---|---|---|---|---|---|---|---|
| ***Gene*** | ***N*** | ***(%)*** | ***Gene*** | ***N*** | ***(%)*** | ***Gene*** | ***N*** | ***(%)*** | ***Gene*** | ***N*** | ***(%)*** |
| *blaOXA-1* | 23 | 8% | *blaCMY-146* | 1 | 0% | *ere(A)* | 1 | 0% | *qnrA1* | 2 | 1% |
| *blaTEM-1A* | 8 | 3% | *blaCMY-2* | 2 | 1% | *mph(A)* | 48 | 17% | *qnrS1* | 5 | 2% |
| *blaTEM-1B* | 82 | 29% | *blaCTX-M-1* | 6 | 2% | *mph(B)* | 3 | 1% | | | |
| *blaTEM-1C* | 8 | 3% | *blaCTX-M-102* | 8 | 3% | | | | | | |
| *blaTEM-1D* | 2 | 1% | *blaCTX-M-14* | 9 | 3% | | | | | | |
| *blaTEM-30* | 1 | 0% | *blaCTX-M-15* | 29 | 10% | | | | | | |
| *blaTEM-34* | 1 | 0% | *blaCTX-M-27* | 1 | 0% | | | | | | |
| *blaTEM-40* | 1 | 0% | *blaCTX-M-3* | 1 | 0% | | | | | | |
| | | | *blaCTX-M-55* | 1 | 0% | | | | | | |
| | | | *blaCTX-M-9* | 2 | 1% | | | | | | |
| | | | *blaSHV-102* | 5 | 2% | | | | | | |
| | | | *blaSHV-12* | 1 | 0% | | | | | | |
| | | | *blaTEM-28* | 1 | 0% | | | | | | |
| | | | *blaTEM-35* | 1 | 0% | | | | | | |
| | | | *blaTEM-52B* | 1 | 0% | | | | | | |

| Aminoglycosides | | | Sulfonamides and Trimetroprim | | | Tetracyclines | | | Other | | |
|---|---|---|---|---|---|---|---|---|---|---|---|
| ***Gene*** | ***N*** | ***(%)*** | ***Gene*** | ***N*** | ***(%)*** | ***Gene*** | ***N*** | ***(%)*** | ***Gene*** | ***N*** | ***(%)*** |
| *aac(3)-Iia* | 1 | 0% | *dfrA1* | 18 | 6% | *tet(A)* | 72 | 26% | *catA1* | 12 | 4% |
| *aac(3)-IIa* | 10 | 4% | *dfrA12* | 6 | 2% | *tet(B)* | 27 | 10% | *strA* | 19 | 7% |
| *aac(3)-Iid* | 4 | 1% | *dfrA14* | 10 | 4% | *tet(D)* | 1 | 0% | *strB* | 10 | 4% |
| *aac(3)-IId* | 7 | 2% | *dfrA17* | 44 | 16% | *tet(J)* | 1 | 0% | *cat* | 1 | 0% |
| *aac(3)-Iva* | 1 | 0% | *dfrA21* | 1 | 0% | *tet(M)* | 1 | 0% | *cmlA1* | 6 | 2% |
| *aac(3)-Via* | 1 | 0% | *dfrA5* | 12 | 4% | *tet(X)* | 1 | 0% | *mdf(A)* | 260 | 93% |
| *aac(6')-Ib-cr* | 12 | 4% | *dfrA7* | 9 | 3% | | | | *floR* | 8 | 3% |
| *aac(6')Ib-cr* | 8 | 3% | *dfrA8* | 2 | 1% | | | | *lnu(F)* | 5 | 2% |
| *aadA1* | 11 | 4% | *sul1* | 64 | 23% | | | | | | |
| *aadA2* | 10 | 4% | *sul2* | 86 | 31% | | | | | | |
| *aadA4* | 1 | 0% | *sul3* | 6 | 2% | | | | | | |
| *aadA5* | 39 | 14% | | | | | | | | | |
| *ant(2")-Ia* | 4 | 1% | | | | | | | | | |
| *ant(3")-Ia* | 22 | 8% | | | | | | | | | |
| *aph(3")-Ib* | 63 | 22% | | | | | | | | | |
| *aph(3')-Ia* | 24 | 9% | | | | | | | | | |
| *aph(3')-Ib* | 1 | 0% | | | | | | | | | |
| *aph(4)-Ia* | 1 | 0% | | | | | | | | | |
| *aph(6)-Id* | 69 | 25% | | | | | | | | | |

In case genes were present twice within a strain, they were only counted once in the resistance gene count.

**S3B Figure.** Acquired resistance gene count among epidemiological subgroups

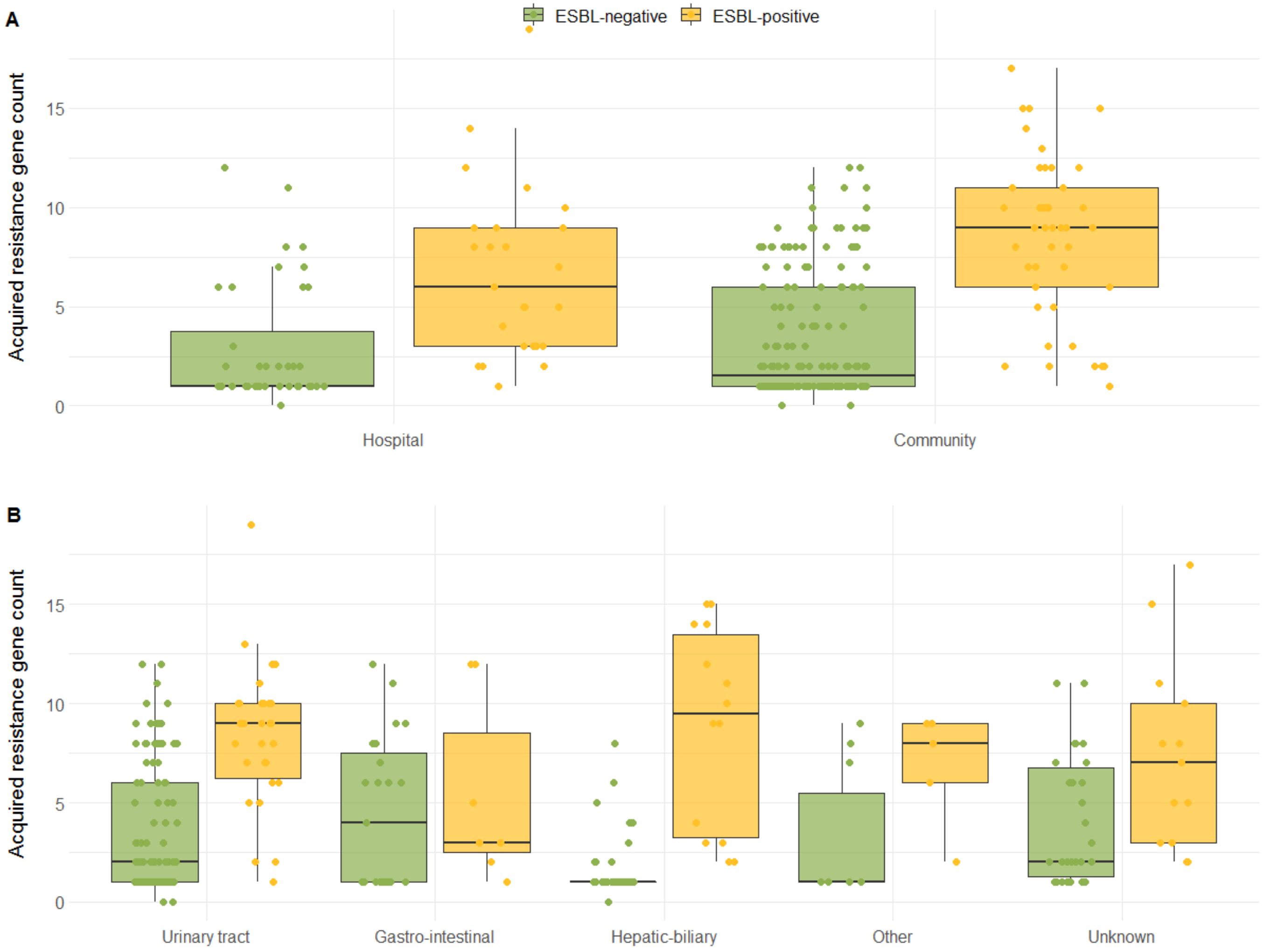


ESBL, extended-spectrum beta-lactamase. ESBL-positivity was based on phenotypic ESBL-production.
Boxplots display median and inter quartile range and every dot represents a single isolate. The ResFinder 3.1.0 database was used to determine acquired resistance genes. **A**. Resistance gene count per onset of infection, stratified for non-ESBL-Ec and ESBL-Ec isolates. **B**. Resistance gene count per primary focus of ECB, stratified for non-ESBL-Ec and ESBL-Ec isolates

**S3C Table.** Pairwise comparisons acquired resistance gene count between epidemiological subgroups

| | Median resistance gene count (IQR) | | | | Pairwise comparisons between groups, within non-ESBL and ESBL[a] | |
|---|---|---|---|---|---|---|
| | Non-ESBL | ESBL | | | Non-ESBL | ESBL |
| **Onset of infection** | | | **Group 1** | **Group 2** | | |
| Community (N = 216) | 2 (1–6) | 9 (6–11) | Community | Hospital | NS[b] | NS |
| Hospital (N = 65) | 1 (1–4) | 6 (3–9) | | | | |
| **Primary focus** | | | **Group 1** | **Group 2** | | |
| Urinary tract (N = 133) | 2 (1–6) | 9 (7–10) | Urinary | GI | NS | NS |
| Hepatic- biliary (N = 60) | 1 (1–1) | 10 (3–14) | Urinary | HB | 2.8e-04*** | NS |
| Gastro-intestinal (N = 30) | 4 (1–8) | 3 (3–9) | Urinary | Other | NS | NS |
| Unknown (N = 43) | 2 (1–7) | 7 (3–10) | Urinary | Unknown | NS | NS |
| Other (N = 15) | 1 (1–6) | 8 (6–9) | GI | HB | 5.8e-03** | NS |
| | | | GI | Other | NS | NS |
| | | | GI | Unknown | NS | NS |
| | | | HB | Other | NS | NS |
| | | | HB | Unknown | 3.1e-05**** | NS |
| | | | Other | Unknown | NS | NS |

ESBL, extended-spectrum beta-lactamase; HB, hepatic-biliary; GI, gastro-intestinal; IQR, interquartile range; NA, not applicable; NS, not significant
[a] Groups were compared with Wilcoxon rank sum Test and *P* values were adjusted with the Holm-Bonferroni correction to adjust for multiple testing. ESBL-positivity was based on phenotypic ESBL-production.
[b] *P* value represents the adjusted *P* value for the comparison of the resistance gene count of Group 1 versus Group 2, within the non-ESBLs or ESBLs (i.e. *P* value 2.8e-04 is the *P* value for the comparison in acquired resistance gene count in urinary versus hepatic-biliary primary focus among non-ESBL *E. coli*)
*, **, *** and **** indicate *P* values ≤0.05, ≤0.01, ≤0.001 and ≤0.0001.
The ResFinder 3.1.0 database was used to determine presence of acquired resistances genes. Gene counts were rounded to whole numbers if applicable.

**S3D Table.** Pairwise comparisons acquired resistance gene count between dominant STs[a]

| | Median resistance gene count (IQR) | | | | Pairwise comparisons between groups, within non-ESBL and ESBL[b] | |
|---|---|---|---|---|---|---|
| | Non-ESBL | ESBL | | | Non-ESBL | ESBL |
| | | | **Group 1** | **Group 2** | | |
| Other ST (N = 150) | 1 (1–6) | 8 (3–12) | ST12 | ST131 | NS | NS |
| ST131 (N = 52) | 2 (1–5) | 9 (6–10) | ST12 | ST38 | NS | NS |
| ST73 (N = 26) | 1 (1–2) | NA | ST12 | ST69 | NS | NS |
| ST69 (N = 21) | 6 (1–8) | 9 (9–9) | ST12 | ST73 | NS | NS |
| ST12 (N = 13) | 1 (1–3) | 3 (3–3) | ST12 | ST95 | NS | NS |
| ST95 (N = 12) | 1 (1–2) | NA | ST131 | ST38 | NS | NS |
| ST38 (N = 7) | 8 (7–8) | 5 (5–8) | ST131 | ST69 | NS | NS |
| | | | ST131 | ST73 | NS | NS |
| | | | ST131 | ST95 | NS | NS |
| | | | ST38 | ST69 | NS | NS |
| | | | ST38 | ST73 | NS | NS |
| | | | ST38 | ST95 | NS | NS |
| | | | ST69 | ST73 | NS | NS |
| | | | ST69 | ST95 | NS | NS |
| | | | ST73 | ST95 | NS | NS |

ESBL, extended-spectrum beta-lactamase; NA, not applicable; NS, not significant; ST, sequence type
ESBL-positivity was based on phenotypic ESBL-production.
[a] Comparisons with category “Other” are not shown; because of heterogeneity in STs this comparison was not considered as informative.
[b] Pairwise comparisons were made with Wilcoxon rank sum Test and *P* values were adjusted with the Holm-Bonferroni correction to adjust for multiple testing.
The ResFinder 3.1.0 database was used for determination of acquired resistance genes. Gene counts were rounded to whole numbers if applicable.
