## Supplementary material for "Extended-spectrum beta-lactamase (ESBL)-producing and non-ESBL-producing *Escherichia coli* isolates causing bacteremia in the Netherlands (2014 – 2016) differ in clonal distribution, antimicrobial resistance gene and virulence gene content": S4 Appendix

**EPIGENEC STUDY - SUPPORTING INFORMATION**

### S4 Appendix - content

**S4A Table.** Detected ExPEC-associated VG per VG category

**S4B Figure.** VG count among epidemiological subgroups

**S4C Table.** Pairwise comparisons VG score between epidemiological subgroups

**S4D Table.** Pairwise comparisons VG score between STs

**S4A Table.** Detected ExPEC-associated VG per VG category[a]

| Adhesins | | Siderophores | | Protectins and invasins | | Toxins | | Other | |
|---|---|---|---|---|---|---|---|---|---|
| **Gene** | **N (%)[b]** | **Gene** | **N (%)[b]** | **Gene** | **N (%)[b]** | **Gene** | **N (%)[b]** | | **N (%)[b]** |
| *yagZ/ecpA* | 271 (96.4) | *sitA* | 233 (82.9) | *ompA* | 235 (83.6) | *usp* | 158 (56.2) | *traT* | 181 (64.4) |
| *fimH* | 266 (94.7) | *fyuA* | 224 (79.7) | *ompT* | 218 (77.6) | *vat* | 101 (35.9) | *malX* | 164 (58.4) |
| *tia* | 124 (44.1) | *chuA* | 158 (56.2) | *kpsM*[c] | 78 (27.8) | *sat* | 91 (32.4) | *iss* | 124 (44.1) |
| *iha* | 111 (39.5) | *iroN* | 135 (48.0) | *tcpC* | 53 (18.9) | *clbB* | 80 (28.5) | *cvaC* | 42 (14.9) |
| *papC* | 103 (36.7) | *iutA* | 32 (11.4) | *ibeA* | 40 (14.2) | *clbN* | 80 (28.5) | *fliC* | 19 (6.8) |
| *papH* | 100 (35.6) | *ireA* | 39 (13.9) | | | *hlyD* | 76 (27.0) | *rfc* | 13 (4.6) |
| *sfa/foc*[c] | 87 (40.1) | | | | | *hlyA* | 72 (25.6) | | |
| *agn43* | 81 (28.8) | | | | | *cnf1* | 66 (23.5) | | |
| *papG* | 57 (20.3) | | | | | *pic* | 45 (16.0) | | |
| *papF* | 55 (19.6) | | | | | *astA* | 29 (10.3) | | |
| *afa/dra*[c] | 43 (15.3) | | | | | *cdtB* | 13 (4.6) | | |
| *nfaE* | 9 (3.2) | | | | | | | | |
| *gafD* | 8 (2.8) | | | | | | | | |
| *bmaE* | 7 (2.5) | | | | | | | | |
| *papE* | 7 (2.5) | | | | | | | | |
| *papA* | 5 (1.8) | | | | | | | | |

VG, virulence genes

[a] The following genes were not detected in any of the isolates: *focE, hra, yfcV* and *tsh* (adhesins) and *hlyF* (toxin)

[b] N indicates numbers of isolates with gene, % of all isolates (N = 281)

[c] The *kpsM*, *afa/dra* and *sfa/foc* operons were considered present if any of the corresponding genes our allelic variants were identified.

**S4B Figure.** ExPEC-associated VG score in different subgroups, stratified for ESBL-positivity[a]

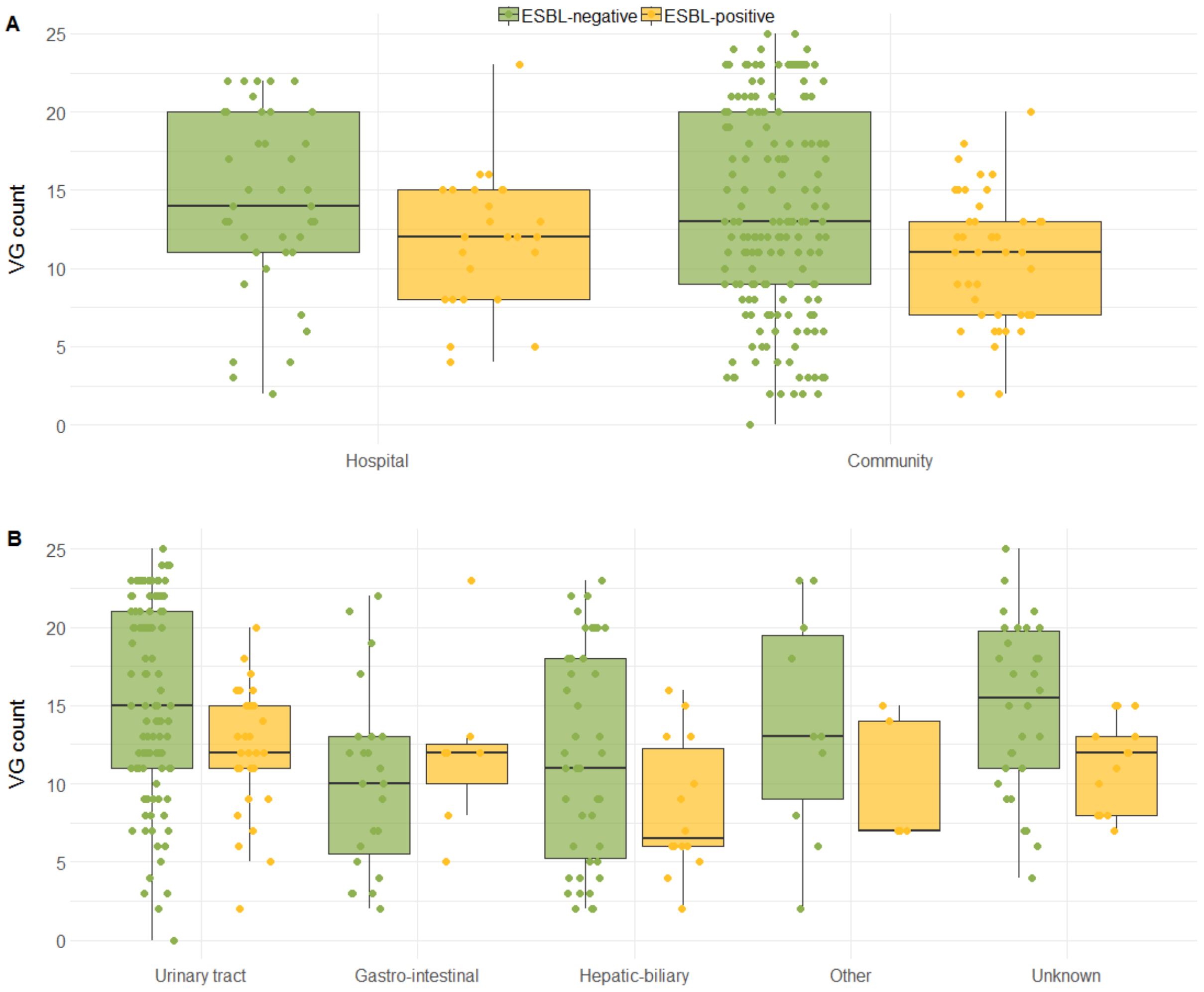


ESBL, extended spectrum beta-lactamase; VG, virulence genes.

[a]ESBL-positivity was based on phenotypic ESBL-production.

Boxplots display median and inter quartile range (IQR) and every dot represents a single isolate. **A**. VG count per onset of infection, stratified for non-ESBL-Ec and ESBL-Ec isolates. **B**. VG count per primary focus of ECB, stratified for non-ESBL-Ec and ESBL-Ec isolates.

**S4C Table.** Pairwise comparisons VG score between epidemiological subgroups

| | Median VG score (IQR) | | Pairwise comparisons between groups, within non-ESBL and ESBL[a] | | | |
|---|---|---|---|---|---|---|
| **Onset of infection** | **Non-ESBL** | **ESBL** | **Group 1** | **Group 2** | **Non-ESBL** | **ESBL** |
| Community (N = 216) | 13 (9–20) | 11 (7–13) | Community | Hospital | NS | NS |
| Hospital (N = 65) | 14 (11–20) | 12 (8–15) | | | | |
| **Primary focus** | | | **Group 1** | **Group 2** | | |
| Urinary tract (N = 133) | 15 (11–21) | 12 (11–15) | Urinary | GI | 0.0072** | NS |
| Hepatic- biliary (N = 60) | 11 (5–18) | 7 (6–13) | Urinary | HB | 0.036* | NS |
| Gastro-intestinal (N = 30) | 10 (5–13) | 12 (8–13) | Urinary | Other | NS | NS |
| Unknown (N = 43) | 16 (11–20) | 12 (8–13) | Urinary | Unknown | NS | NS |
| Other (N = 15) | 13 (8–20) | 7 (7-14) | GI | HB | NS | NS |
| | | | GI | Other | NS | NS |
| | | | GI | Unknown | NS | NS |
| | | | HB | Other | NS | NS |
| | | | HB | Unknown | NS | NS |
| | | | Other | Unknown | NS | NS |
| **Urinary catheter** | | | **Group 1** | **Group 2** | | |
| No (N = 184) | 13 (9-20) | 10 (7-13) | No catheter | Catheter | NS | NS |
| Yes (N = 97) | 13 (7-18) | 13 (11-15) | | | | |
| **30-day mortality** | | | **Group 1** | **Group 2** | | |
| Alive (N = 238) | 13 (9-20) | 12 (8-15) | Alive | Deceased | NS | NS |
| Deceased (N = 43) | 12 (6-18) | 11 (7-14) | | | | |
| **Admission ward** | | | **Group 1** | **Group 2** | | |
| Non-ICU (N = 240) | 13 (9-20) | 12 (7-15) | Non-ICU | ICU | NS | NS |
| ICU (N =41) | 13 (6-18) | 12 (10-14) | | | | |

ESBL, expended-spectrum beta-lactamase; HB, hepatic-biliary; GI, gastro-intestinal; IQR, interquartile range; NA, not applicable; NS, not significant; VG, virulence gene.
[a] Pairwise comparisons were made with Wilcoxon rank sum Test and *P* values were adjusted with the Holm-Bonferroni correction to adjust for multiple testing.
* and ** indicate *P* values ≤0.05 and ≤0.01. Gene counts were rounded to whole numbers if applicable. ESBL-positivity was based on phenotypic ESBL-production.

**S14D Table.** Pairwise comparisons VG score between dominant STs[a]

| | Median VG score (IQR) | | Pairwise comparisons between groups within non-ESBL and ESBL[b] | | | |
|---|---|---|---|---|---|---|
| | **Non-ESBL** | **ESBL** | **Group 1** | **Group 2** | **Non-ESBL** | **ESBL** |
| Other ST (N = 150) | 11 (7–17) | 8 (6 –11) | ST12 | ST131 | 3.2e-05**** | NS |
| ST131 (N = 52) | 13 (12–15) | 13 (12–15) | ST12 | ST38 | NS | NS |
| ST73 (N = 26) | 22 (20–23) | - | ST12 | ST69 | 5.5e-05**** | NS |
| ST69 (N = 21) | 11 (9–12) | 8 (7–8) | ST12 | ST73 | NS | - |
| ST12 (N = 13) | 22 (21–23) | 23 (23–23) | ST12 | ST95 | 0.032* | - |
| ST95 (N = 12) | 18 (17–19) | - | ST131 | ST38 | NS | NS |
| ST38 (N = 7) | 7 (6–7) | 8 (7–8) | ST131 | ST69 | 4.4e-03** | NS |
| | | | ST131 | ST73 | 1.9e-07**** | - |
| | | | ST131 | ST95 | 41.9e-03** | - |
| | | | ST38 | ST69 | NS | NS |
| | | | ST38 | ST73 | NS | - |
| | | | ST38 | ST95 | NS | - |
| | | | ST69 | ST73 | 2.0e-07**** | - |
| | | | ST69 | ST95 | 5.8e-05**** | - |
| | | | ST73 | ST95 | 0.032* | - |

ESBL, extended-spectrum beta-lactamase; NS, not significant; ST, sequence type; VG, virulence gene.
[a] Comparisons with category "Other" are not shown; because of heterogeneity in STs this comparison is not considered as informative.
[b] Groups were compared with Wilcoxon rank sum Test and P-values were adjusted with the Holm-Bonferroni correction to adjust for multiple testing. ESBL-positivity was based on phenotypic ESBL-production.
*, **, *** and **** indicate P-values ≤0.05, ≤0.01, ≤0.001 and ≤0.0001.
Gene counts were rounded to whole numbers if applicable.
